# Perivascular Adipose Tissue: Quantitative analysis by morphometry and stereology in rodents

**DOI:** 10.1101/820415

**Authors:** Felipe Demani Carneiro, Stephanie Christinne Sinder Mello, Emiliana Barbosa Marques, Rogerio Barbosa Magalhaes Barros, Christianne Bretas Vieira Scaramello, Caroline Fernandes-Santos

## Abstract

The perivascular adipose tissue (PVAT) provides mechanical support to blood vessels and modulates vascular physiology in obesity. Our goal is to provide a reproductive protocol using morphometric and stereological tools to assess PVAT morphology. The thoracic aorta from male Wistar rats (n=6) and C57BL/6 mice (n=7) underwent routine histological procedures, and two independent observers analyzed the same set of digital images. Agreement and reproducibility were assessed. Both observers showed that the diameter of rat brown adipocytes is larger than mice (P<0.002) as expected, and that the number density (Q_A_) of brown adipocytes is smaller in rats compared to mice (P<0.01). Considering lipid droplets, observer #1 reported that in rats they were larger (P<0.005) and had a higher volume density (V_V_) than mice (P=0.035), but observer #2 found the opposite for lipid droplet diameter (P=0.001). White adipocytes were not found in the PVAT. Bland-Altman plots demonstrated agreement and reproducibility between observers since the means are close to the main difference (bias) and within the 95% limits of agreement. In conclusion, the methodology proposed can quantify morphological aspects of the aorta PVAT in rodents. It is reproducible and can be performed by both expert and inexperienced researchers, once they know how to recognize the structures of interest to be measured.

## INTRODUCTION

Virtually all arteries, except brain arteries, are surrounded by a significant amount of perivascular adipose tissue (PVAT) (1). It was thought that the PVAT was only responsible for the mechanical protection of vessels against neighboring tissues during contraction (2). However, recent studies have shown that the PVAT is responsible for the mechanical support of blood vessels and the secretion of various substances. Among them, there are a large number of metabolically active adipokines, chemokines (e.g., interleukin-6 and tumor necrosis factor alpha), hormone-like factors (e.g., leptin, adiponectin, and resistin), and vasoactive substances (e.g., prostacyclin, adiponectin and prostaglandin) (3).

In rodents, the thoracic aorta PVAT consists of brown adipocytes that morphologically resemble the classic brown adipocytes found in interscapular brown adipose tissue (iBAT). The abdominal aorta PVAT is composed of a mixture of brown and white adipocytes, and the PVAT from other arteries such as mesentery, femoral and carotid arteries consists only of white adipocytes (4). In conditions such as obesity and diabetes, the PVAT becomes dysfunctional. It expands in size, accumulates inflammatory cells, and changes its secretory profile of several adipokines and proinflammatory cytokines (5).

Morphometry is a two-dimensional quantitative method, which aims to determine parameters such as lengths, perimeters, and areas. It can be easily performed using an appropriate image analysis software such as ImageJ (free, https://imagej.nih.gov/ij/?) and Image Pro Plus (paid, http://www.mediacy.com/imageproplus). On the other hand, design-based stereology methods rely on statistical sampling principles and stochastic geometric theory to estimate quantitative parameters of three-dimension geometric objects in complex tissue structures (6, 7). Stereology uses test-system probes such as points, lines, and frames to estimate volumes, surfaces, lengths and numbers of the structure of interest. Stereology uses tissue sections that only show two-dimensional information and provides two- and three-dimensional information, whereas morphometry only makes assumptions in two-dimensions.

It is important to assess the morphological characteristics of PVAT by morphometric tools since this tissue has an active influence on vascular physiology and is considered a distinct tissue regarding anatomical location, histological characteristics and molecular biology (3). To date, there is no methodology published describing how to assess morphometric and stereological parameters of PVAT. Thus, our purpose is to provide some morphometric tools to study PVAT morphology. For this, we used the thoracic aorta PVAT of albino Wistar rats and C57BL/6 mice. Since rat and mice differ in body size, we expect that rat PVAT cells are larger than mice, and thus we tested whether the methodology proposed can detect differences between rats and mice. Also, the analyses were performed by two independent observers with and without previous experience in morphological quantification, to assess the impact of expertise on reproductivity.

## MATERIALS AND METHODS

### PVAT collection and processing

The handling and experimental protocols were approved by the local Ethics Committee to Care and Use of Laboratory Animals (CEUA#647/15). The study was performed in agreement with the Animal Research Reporting in Vivo Experiments ARRIVE guidelines and the Guideline for the Care and Use of Laboratory Animals (US NIH Publication N° 85-23. Revised 1996) (8). Male C57BL/6 mice (n=7) and male albino Wistar rats (n=6) with five months old were used. Animals were obtained from colonies maintained at the Federal Fluminense University Animal Care Facility and kept under standard conditions (12h light/dark cycles, 21±2°C, humidity 60±10% and air exhaustion cycle 15min/h). Food and water were offered *ad libitum*, and the body mass was measured at the time of euthanasia. Average body mass was 433±21.4g for rats and 28±1.6g for mice.

For tissue collection, animals were submitted to six hours fasting and were deeply anesthetized with ketamine 100.0 mg/kg (Francotar^®^, Virbac, Brazil) and xylazine 10.0 mg/kg ip (Virbaxyl 2%^®^, Virbac, Brazil). The thoracic aorta was dissected, its proximal segment close to the aortic arc was immersed in Millong formalin (4% w/v in 0.1M phosphate buffer pH 7.2) for 48 hours, and then it followed the routine histological processing and embedding (Paraplast Plus, Sigma–Aldrich, St. Louis, MO, USA). Nonconsecutive sections were obtained to avoid counting the same structures (30 µm distance) and then stained with hematoxylin and eosin. Digital images of the PVAT were obtained using a Leica DM750 microscope (Wetzlar, German) coupled to a video camera Leica ICC50 HD (Wetzlar, German).

### PVAT morphometry

Morphometry was performed in the computer-based software Image Pro^®^ Plus v. 5.0 (Media Cybernetics, Silver Spring, MD, USA), which allows the counting, measurement, and classification of objects. Ten nonconsecutive images of the PVAT were acquired per animal (image resolution 2,048 × 1,536 pixels .jpeg). Brown adipocytes have a polygonal shape and thus we measured the largest and smallest diameter of 10 cells per image using the line feature, summing 200 measurements from 100 adipocytes (Fig. 1a). Since lipid droplets have a spherical shape, we used the circle feature on Image Pro Plus to measure their average diameter. Two random adipocytes per image had all their lipid droplets measured and thus 20 adipocytes were analyzed (Fig. 1b). Aorta wall was not evaluated in the present study since this methodology has been described elsewhere (9).

**Figure 1.**
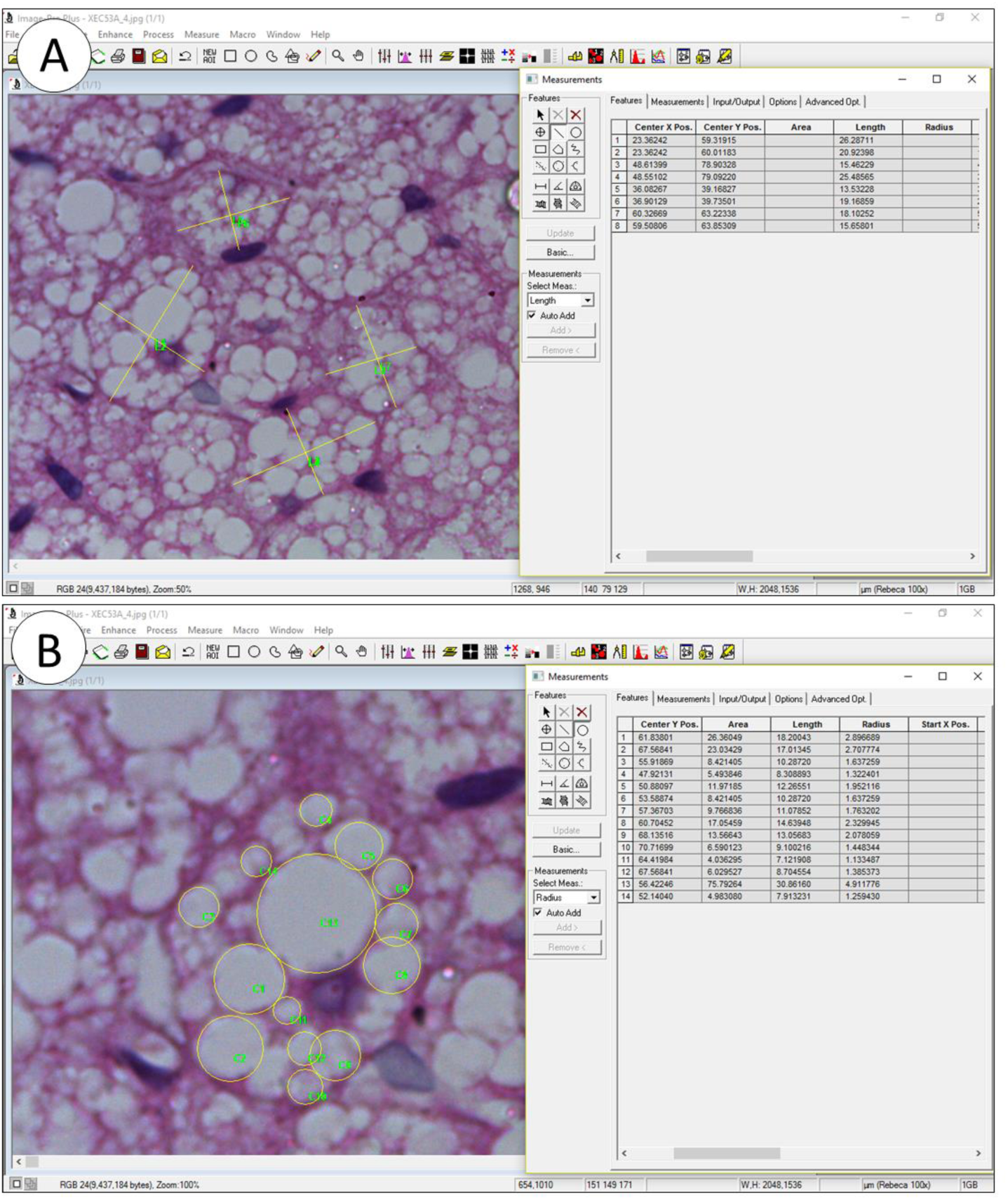
Morphometry performed on Image Pro Plus to assess diameter. A, brown adipocytes had their biggest and smallest diameter measured by the line feature. B, lipid droplets had their diameter measured by the circle feature. Images in A and B are the same, but the zoom in tool was used to allow a better visualization of the structures.

### PVAT stereology

Stereology was performed in STEPanizer (http://www.stepanizer.com/), that is a free easy-to-use computer-based software tool for stereological assessment of digitally captured images (10). It creates test systems that are superimposed on digital images. Images can be scaled, and it has a counting module and an export function of data to spreadsheet programs. Monitor resolution was 768 × 1,024 and images used for stereology were the same used for morphometry. The number density (*Q*_*A*_) of brown and paucilocular adipocytes was estimated in a test-frame of 2,966 µm^2^ and a guard area of 150 pixels width. All adipocytes inside the test-frame were considered but not the ones that hit the forbidden line or its extensions to avoid overestimation. The *Q*_*A*_ was calculated as the number of cells inside the test-frame divided by test-frame area, expressed in mm^2^. Brown adipocytes were considered as the multilocular cells possessing lipid droplets of varied sizes and a central nucleus (when visible). Paucilocular adipocytes are cells with intermediate morphology between that of white and brown adipocytes, and they were considered as the multilocular cells possessing a single and pronounced central lipid droplet surrounded by small lipid droplets and a peripherical nucleus, when visible (11). White adipocyte where not present.

The volume density (*V*_*V*_) of lipid droplets was estimated by point counting in a 49-points test system and a guard area of 10 pixels width. The *V*_*V[lipid droplet]*_ was estimated as *P*_*P[lipid droplet]*_*/ P*_*T*_, where *P*_*P*_ is the number of points that hit lipid droplets and *P*_*T*_ is the total test points, in this case, 49 (Fig. 2b). In general, counting 100-200 points per study subject is considered as a sufficient count when the structure of interest is representative, and to do additional sampling and counting would be inefficient because biological variation cannot be altered by more sampling and counting (12-14). In our experience, lipid droplets occupy about ⅓ to ¼ of the tissue section and considering that we analyzed 10 sections, a total of 490 points were used to estimate *V*_*V[lipid droplets]*_, which would be appropriate.

**Figure 2.**
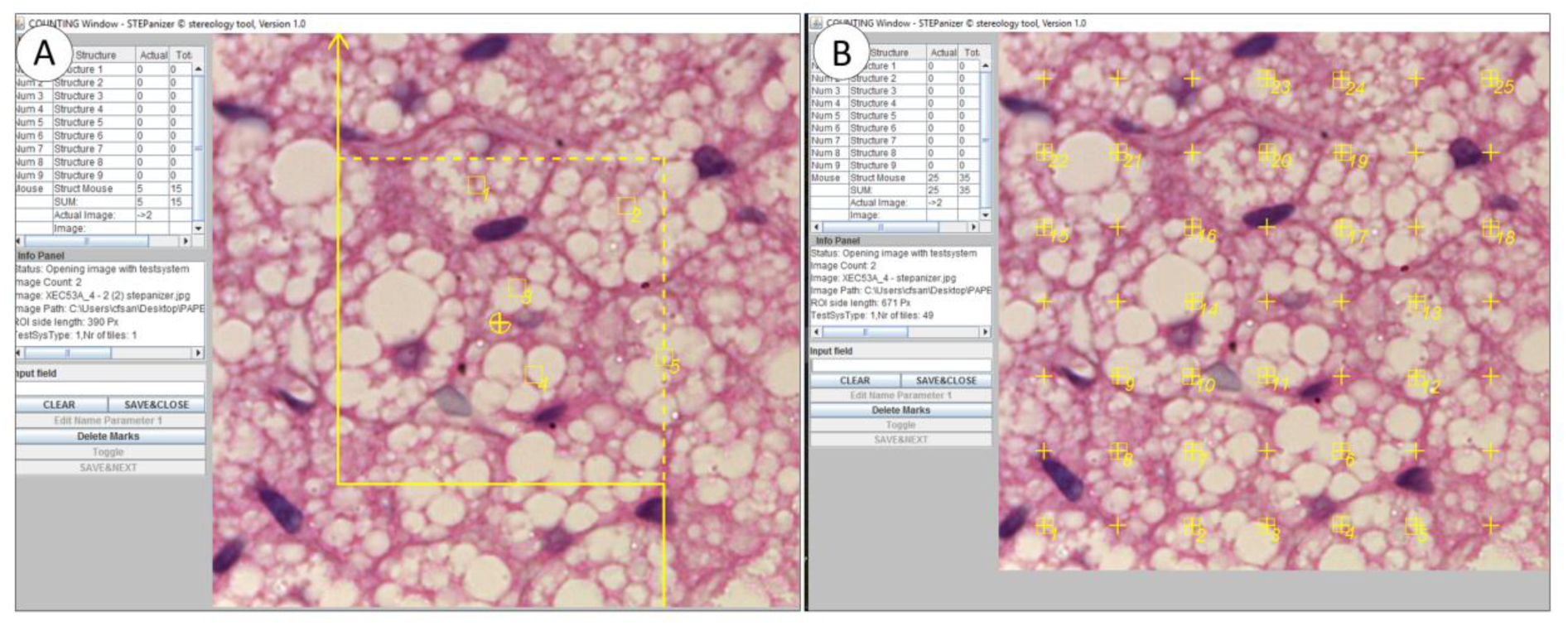
Stereology performed on STEPanizer to assessed number and volume densities. A, the number density (Q_A_) of brown adipocytes is estimated by counting the number of profiles inside the counting frame (in this example, n=5). Cells that touch the forbidden line are not considered to avoid overestimation (continuous line). B, volume density (V_V_) of lipid droplets was estimated by point counting in a 49-points test system. Points that hits lipid droplet profiles are considered (in this example, n=25).

### Observers

All measurements were performed by two blind observers (#1 and #2), that used the same set of images, and analyzed them using their own computer, to assess the reproducibility of the method. Observer #1 had a previous experience performing morphological quantification of PVAT in Image Pro Plus and STEPanizer softwares, but observer #2 had no previous experience about morphological tools and softwares of tissue quantification. Before quantifying, a third researcher (senior advisor) helped the two observers to perform system calibration and discussed with them how to identify, count and measure the structures of interest.

### Statistics analysis

Data are expressed as mean ± standard deviation, and it was tested for normality and homoscedasticity of variances. Rat and mice parameters obtained from the same observer were compared with the Mann-Whitney U test. This test was also used to compare a similar parameter obtained by observers #1 and #2. Bland-Altman graphs were created to assess the agreement between the two observers. The difference between observer #2 and #1 (Obs2 - Obs1) was plotted against the average of each parameter. The bias of one observer to the other is represented by the mean of the differences and the 95% limits of agreement (mean ± 2S.D.). Graph Pad® Prism v.6.0 (La Jolla, CA, USA) was used to perform all analysis and a *P*<0.05 was considered statistically significant.

## RESULTS

### PVAT quantification

Table 1 shows the morphometric and stereological parameters obtained by observers #1 and #2 using the same set of images acquired from mice and rat PVAT. For morphometry, observers #1 and #2 found that brown adipocytes of rats are larger than mice (+26% *P*=0.0023, +18% *P*=0.0047, respectively). However, whereas observer #1 found lipid droplets larger in rats compared to mice (+16% *P*=0.0047), observer #2 found the opposite (−19% *P*=0.0012). This difference between observers seems to be related to the quantification of rat lipid droplet diameter since average mice lipid droplet diameter was similar between observers.

**Table 1.**
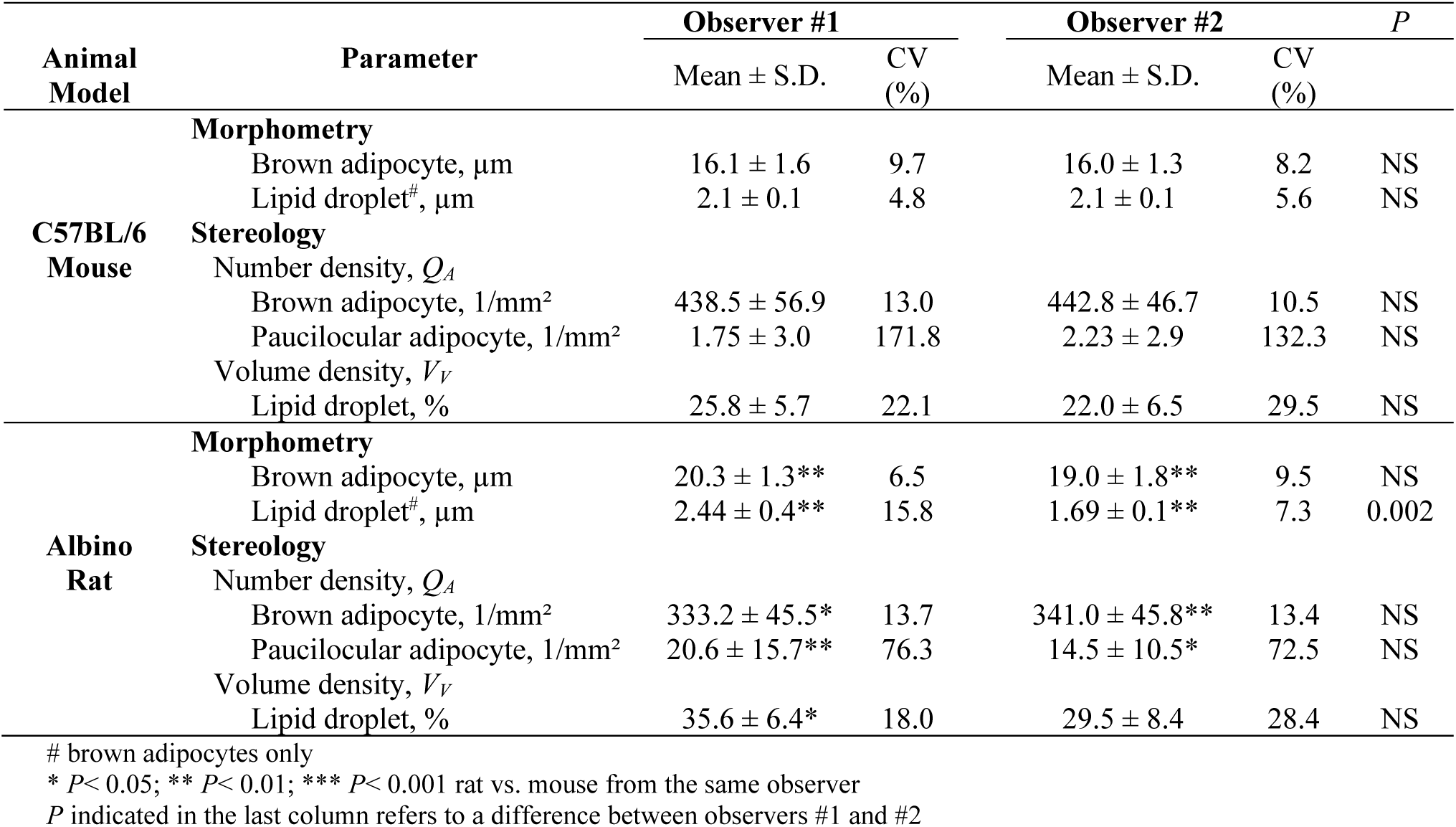
Comparison between observer #1 and observer #2

For stereology, both observers detected a smaller number density of brown adipocytes in rats compared to mice (+24% *P*=0.0134, +23% *P*=0.0041, respectively), and the same is true for paucilocular adipocytes (+1,077% *P*=0.0052, +550% *P*=0.0256, respectively). Observer #1 showed a higher volume density of lipid droplets in rats (+38% *P*=0.035), but observer #2 found only a small trend with no significance. It is important to notice that the coefficient of variation is low for most parameters analyzed, but not for *Q*_*A[paucilocular adipocytes]*_ and *V*_*V[lipid droplets]*._

Overall morphological findings are summarized in Fig 3, where rats display larger brown adipocytes with larger lipid droplets compared to mice. Therefore, in rats, fewer cells are found per area (*Q*_*A[brown adipocyte]*_ rat < mice) and lipid droplets occupy a bigger volume within tissue section (*V*_*V[lipid droplets]*_ rat > mice).

**Figure 3.**
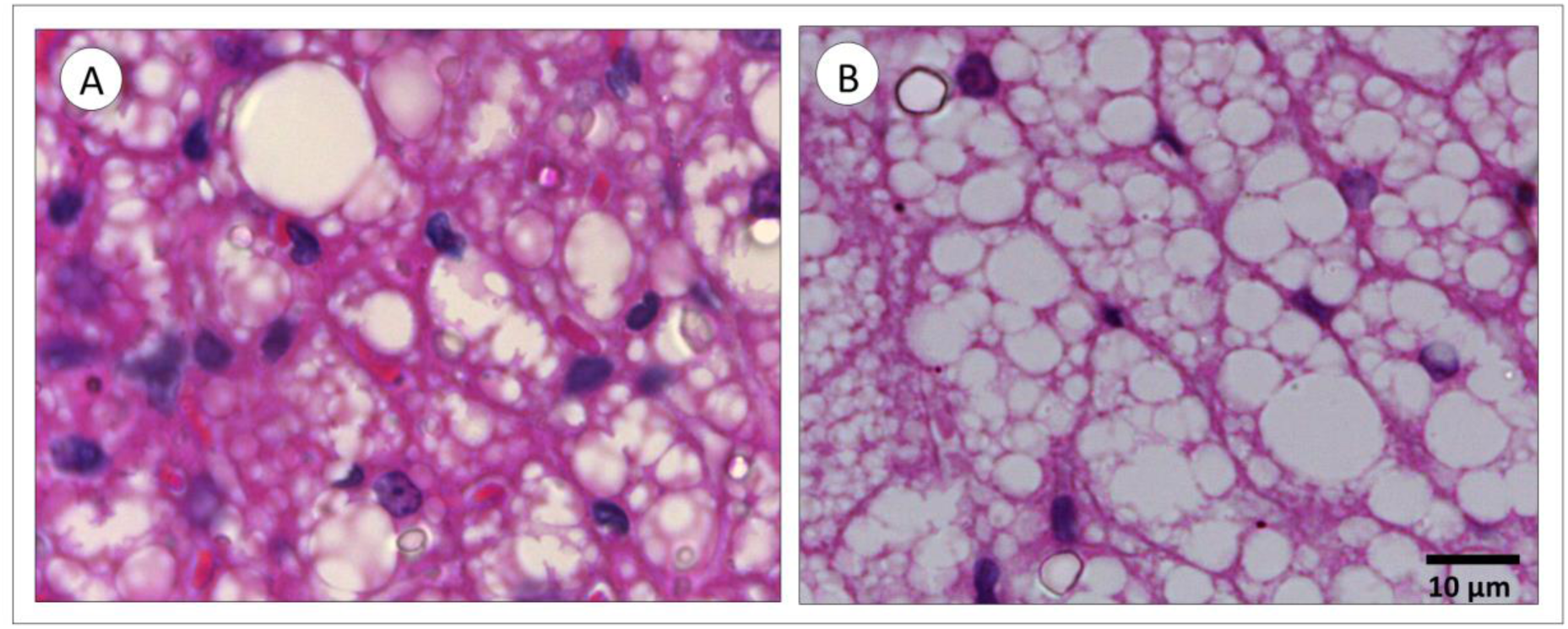
Perivascular adipose tissue from mice (A) and rat (B). In rats, brown adipocytes and lipid droplets are larger than mice. As a consequence, the number density (Q_A_) of brown adipocytes in rats is smaller, and the volume density (V_V_) of lipid droplets is bigger than mice.

### Inter-observer agreement and reproducibility

Bland-Altman test parameters are shown in Table 2. Curves in Figs 4-5 have plotted the average of mice plus rat parameters (x-axis) against the difference between observer #1 and #2 (y-axis). An overall analysis of the five curves shows that morphometric (Fig 4) and stereological (Fig 5) quantification agreed among observers. However, in Fig 4b the data close to 0.0 regarding obs2-obs1 are from rats, whereas all mice data are negatives, which increases the bias and the 95% confidence interval. When analyzed independently, the average lipid droplet diameter bias (±S.D) for rats is 0.046±0.06 and for mice −0.624±0.46 (data not shown). Also, most positive data plotted in Fig 5a are from mice, and thus observer #2 found values bigger than observer #1 for the same animal.

**Table 2.** Reproducibility between observers evaluated by the Bland-Altman test

| Parameter |  |  | Limits of agreement |  |
| --- | --- | --- | --- | --- |
|  | Bias | SD Bias | Lower (CI) | Upper (CI) |
| **Morphometry** |  |  |  |  |
| Brown adipocyte, $\mu\text{m}$ | -0.6723 | 1.259 | -3.140 | 1.795 |
| Lipid droplet<sup>#</sup>, $\mu\text{m}$ | -0.3152 | 0.476 | -1.248 | 0.618 |
| **Stereology** |  |  |  |  |
| Number density, $Q_A$ | | | | |
| Brown adipocyte, $1/\text{mm}^2$ | 5.900 | 11.60 | -16.84 | 28.64 |
| Paucilocular adipocyte, $1/\text{mm}^2$ | -2.566 | 6.968 | -16.22 | 11.09 |
| Volume density, $V_V$ | | | | |
| Lipid droplet, % | -4.908 | 3.329 | -11.43 | 1.616 |

CI, 95% confidence interval
<sup>#</sup> only from brown adipocytes

**Figure 4.**
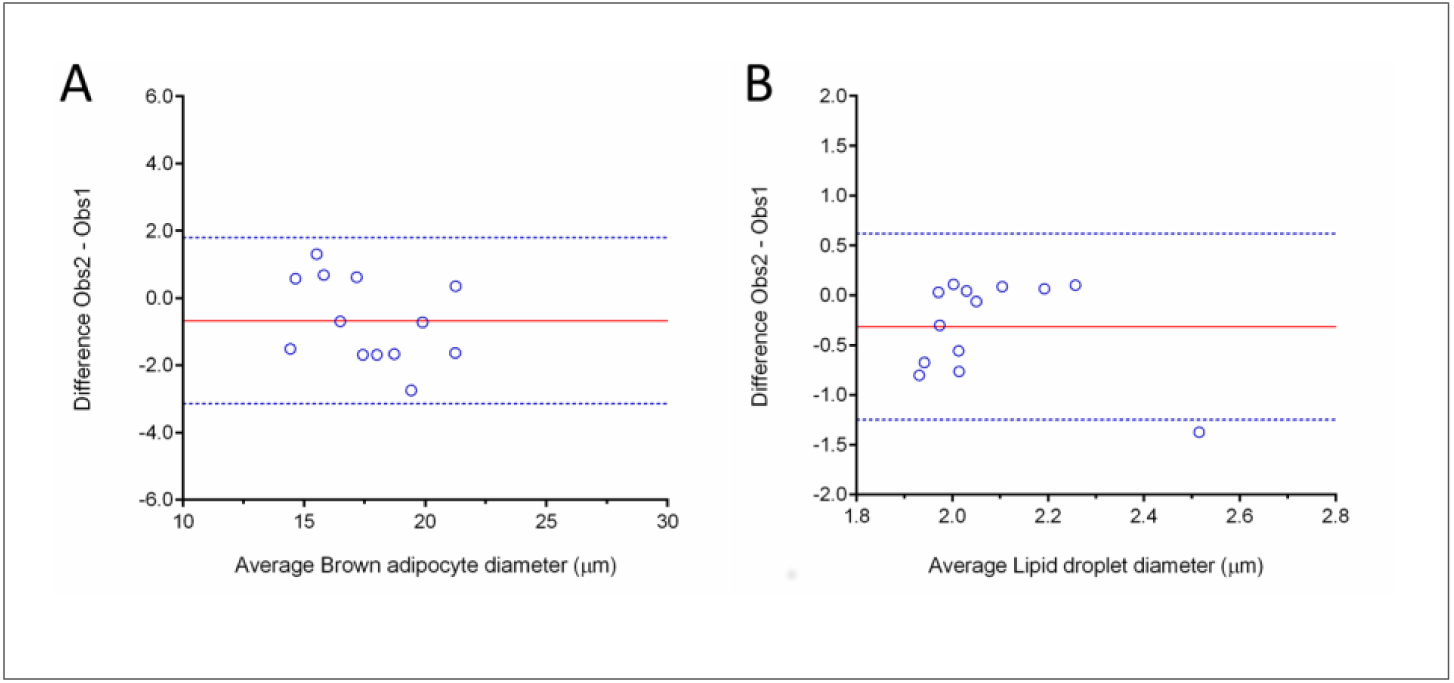
Morphometry reproductivity between observers assessed by the Bland-Altman method. Graphs plot the difference between observers (Obs2-Obs1) against the mean ([Obs1+Obs2]/2). Brown adipocyte diameter is shown in **A** and lipid droplet diameter in **B**.

**Figure 5.**
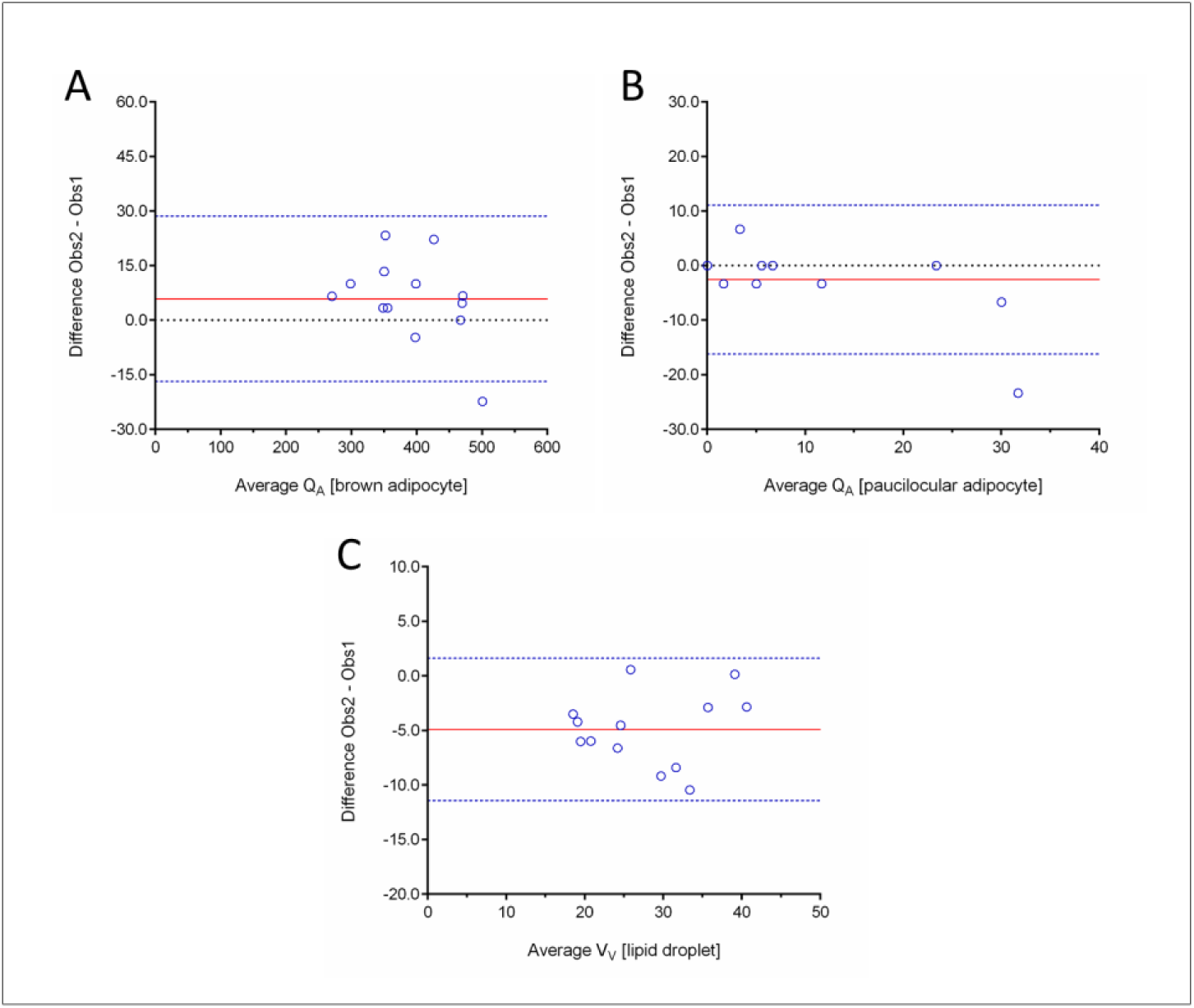
Stereology reproductivity between observers assessed by the Bland-Altman method. Graphs plot the difference between observers (Obs2-Obs1) against the mean ([Obs1+Obs2]/2). **A**, number density (Q_A_) of brown adipocytes. **B**, Q_A_ of paucilocular adipocytes. **C**, volume density (V_V_) of lipid droplets (brown adipocytes only).

## DISCUSSION

We provided some morphometric and stereological tools to allow the study of PVAT morphology in rodents. Morphometry is a quantitative method that can be used to determine lengths, perimeters and areas of biological specimens in two-dimension, whereas stereology estimates volumes, surfaces, lengths and numbers of structures in tissue sections that only provides two-dimensional information, providing two-and three-dimensional information. The methodology presented can detect morphological differences among the PVAT of albino Wistar rats and C57BL/6 mice, such as the average size of brown adipocytes and their lipid content. These parameters are extremely important since they can be used to understand PVAT remodeling in obesity by using animal models such as diet-induced obesity and genetic models of obesity. Assessing aorta and its PVAT remodeling, together with gene expression and functional approaches will allow the understand of mechanisms underlying arterial dysfunction in obesity. This integrated technique approach to exploit vascular biology will help to elucidate the onset and development of vascular dysfunction in obesity.

Morphometry and stereology are considered distinct quantitative methods. They possess several peculiarities, different purposes and are executed by specific image analysis software. Despite their inherent methodological differences, the data generated have good reproducibility. These methodologies allow inter-group comparisons, and their theoretical background is well established and accepted. Of note, only a small training is required so that inexperienced researchers not habituated with the methodology can easily perform it (6). In the present study, morphometry and stereology data obtained by the two observers agreed, which ratifies that even inexperienced researchers can perform the methodology and obtain reproductive data.

Observers #1 and #2 did not have difficult to execute the methodologies proposed, but observer #2 (inexperienced) reported some difficult to identify lipid droplet boundary in the images provided. It might justify the inter-observer difference found for rat lipid droplet diameter, and an absence of difference in the volume density of lipid droplets between rats and mice for observer #2. Digital images were obtained using the 100x objective of an optical microscope, that is the highest magnification possible in this system. An alternative to lipid droplet quantification might be the use of electron photomicrography since they provide a higher magnification, allowing a better visualization of tissue organelles and their boundaries. However, it is an expensive technique, and not all laboratories perform it as a routine. Despite this limitation, the Bland-Altman data reported agreement and reproducibility between observers.

The body has two types of adipose tissues, the white (WAT) and brown adipose tissue (BAT). WAT is important for energy storage, and it is capable of rapidly increasing its size by adipocyte hypertrophy and hyperplasia. WAT depots are found surrounding internal organs (visceral fat) and under the skin (subcutaneous fat), and visceral fat expansion in obesity is associated with increased cardiovascular risk (15, 16). BAT is responsible for adaptive thermogenesis, being consistently identified in adult humans in the cervical-supraclavicular, perirenal, adrenal, paravertebral and surrounding large vessels as PVAT (15). In some individuals, brown adipocytes are also found within the WAT depot and are referred to as beige adipocytes (17, 18). Since brown adipocytes burn fat, several strategies are under investigation to increase the number and activity of beige cells as an attempt to induce weight loss, improve metabolism, and reduce cardiovascular risk (19-21).

White adipocytes are spherical cells with ∼90% of their volume comprising a single cytoplasmic lipid droplet and a peripheric nucleus, whereas brown adipocytes are polygonal cells with a roundish nucleus and have several cytoplasmic lipid droplets (11, 19, 22). The paucilocular adipocyte is considered as an intermediate step of white-to-brown adipocyte transdifferentiation. Consequently, it presents an intermediate morphology between white and brown adipocytes, exhibiting a large vacuole surrounded by at least five small lipid droplets (11, 23, 24). Paucilocular adipocytes are found in all adipose tissue deposits in humans and rodents (25). In our study, we noticed paucilocular adipocytes in rats and mice thoracic aorta PVAT, but not in all animals, and they were more often seen in rats compared to mice. The high coefficient of variance for *Q*_*A[paucilocular adipocyte]*_ indicates that the paucilocular adipocyte is not a frequent cell in the thoracic aorta PVAT of male albino Wistar rats and C57BL/6 mice. Thus, if the researcher has the aim to evaluate this subtype of adipocyte, more tissue sections are necessary to obtain an unbiased estimation of its numerical density.

## CONCLUSIONS

In conclusion, the methodology proposed can quantify morphological aspects of the aorta PVAT in rodents. It is reproducible and can be performed by both expert and inexperienced researchers, once they know how to recognize the structures of interest to be measured.

## ACKNOWLEDGEMENTS

Authors are thankful for Dilliane da Paixão Rodrigues Almeida for her technical assistance.

## AUTHORSHIP

Fernandes-Santos C conceived and designed the experiments; Carneiro FD and Mello SCS performed the experiments; Fernandes-Santos C analyzed and interpreted the data; Marques EB, Barros RBM, and Scaramello CBV contributed with reagents, materials, and animals; Carneiro FD and Fernandes-Santos C wrote the paper

